# In-silico modelling of the mitogen-activated kinase (MAPK) pathway in colorectal cancer: mutations and targeted therapy

**DOI:** 10.1101/2023.04.18.537359

**Authors:** Sara Sommariva, Silvia Berra, Giorgia Biddau, Giacomo Caviglia, Federico Benvenuto, Michele Piana

## Abstract

Chemical reaction networks are powerful tools for computing the complex nature of cancer’s onset, progression, and therapy. The main reason for their effectiveness is in the fact that these networks can be rather naturally encoded as a dynamical system whose asymptotic solution mimics the proteins’ concentration profile at equilibrium. The paper relies on this mathematical approach to investigate global and local effects on the chemical reaction network of the colorectal cancer, triggered by partial and complete mutations occurring in its mitogen-activated kinase (MAPK) pathway. Further, this same approach allowed the in-silico modelling and dosage of a multi-target therapeutic intervention that utilizes MAPK as its molecular target.

## 1 INTRODUCTION

For many years, the primary sources of anti-cancer intervention just consisted in chemiotherapy, where specific drugs are used to kill cancer cells, and surgery. However, surgery is not always feasible, while the broad-spectrum of chemioterapy drugs, which attack indiscriminately cancer and fast-growing healthy cells, often results in high toxicity (Lowenthal and Eaton, 1996). To overcome this low specificity of chemiotherapy, novel biology-based approaches have been introduced in routine cancer therapies. In particular, following the advances of human genome sequencing, in the last twenty years an increasing number of molecular targeted drugs have been introduced. This kind of therapeutic intervention aims at slowing down cancer progression and metastasis by targeting specific molecules somehow involved in the genetic alterations that underlie cancer onset (Lee et al., 2018; Bedard et al., 2020; Zhong et al., 2021). Additionally, synergies of multiple targeted drugs combined in a single therapy have been investigated to reduce resistance to single-agent therapies (Jin et al., 2023). Identifying novel molecular targets and optimizing dosage and combination of the corresponding drugs remains a challenging problem, where the number of possible therapies to be tested vastly exceeds clinical resources, in terms of both financial and time resources. In this scenario, systems biology models could play a crucial role in identifying the most promising candidates for clinical trials and in elucidating the molecular mechanisms underlying targeted drugs synergies (Chen et al., 2015; Rocca and Kholodenko, 2021).

At a worldwide level, colorectal cancer (CRC) is the third most frequent cancer in male population and the second one among women (Xi and Xu, 2021; Sung et al., 2021; Biller and Schrag, 2021). Screening programs have contributed in reducing the incidence of later-onset cases while an alarming increase in early-onset cases and in corresponding CRC-related mortality among younger people have been observed (Saad El Din et al., 2020; Wu and Lui, 2022; Sinicrope, 2022). At a molecular level CRCs are highly heterogeneous pathologies with differences across age groups. It has been estimated that five to ten tumor-specific driver mutations usually concur in individual cancers and that the most frequent alterations in CRC pertain to TP53, APC, KRAS, PTEN, SMAD4, PIK3CA, BRAF, and AKT (Tariq and Ghias, 2016; Tortolina et al., 2015; Anderson et al., 2019). Among these genes, KRAS, APC, SMAD4, and TP53 belong to four different pathways, namely MAPK, WNT, TGF*β*, and TP532, each one acting at a different functional stage of cell development, ranging from stem cell renewal to cell growth, division and apoptosis (Armaghany et al., 2012). The first targeted therapies for CRCs, namely cetuximab (Jonker et al., 2007), which inhibits the epidermal growth factor receptor (EGFR), and bevacizumab (Los et al., 2007; Rosen et al., 2017), against the vascular endothelial growth factor A (VEGF-A), were approved by the Food and Drug Administration (FDA) in 2004. Since then, various pathways have been proved to offer ideal sites for targeted therapies, and an increasing number of novel agents have been developed (for a comprehensive review we refer to (Tiwari et al., 2018), (Xie et al., 2020), and references therein). However, to date only a few CRC-related pathways have been successfully inferred due to the complex signaling network that makes it hard to completely inhibit specific biological interactions. As a consequence, many proposed therapies have not passed the preclinical status or the phase I trial, highlighting the need for systems biology models capable of guiding the choice of which drugs to test so as to avoid waste of resources.

It is clear that a mathematical model aiming at capturing the complex nature of CRC onset, progression, and therapy cannot consider altered and targeted proteins and corresponding pathways in isolation but must integrate them within proper chemical reaction networks (CRNs) (Tortolina et al., 2015). An extensive CRN for CRC has been recently introduced for modelling signal transduction during the G1-S transition phase in colorectal cells (Tortolina et al., 2015; Castagnino et al., 2016). Such a CRN, henceforth denoted as CR-CRN, comprises 10 different pathways, including all four previously mentioned ones, for a total of 419 proteins interacting in 850 chemical reactions. Following standard mathematical procedures based on the law of mass action (Feinberg, 1987; Chellaboina et al., 2009; Yu and Craciun, 2018), the CR-CRN has been mapped into a system of 419 autonomous ordinary differential equations (ODEs) whose solutions describe the behaviour of protein concentrations, which evolve in time until the network reaches a globally asymptotically stable state that also represents an equilibrium of the system (Ingalls, 2013; Sommariva et al., 2021a). It has been conjectured that the CR-CRN satisfies the so called global stability condition (Sommariva et al., 2021a), meaning that a unique stable state exists once fixed the initial values of the protein concentrations or, more precisely, once fixed the total moiety within the conservation laws of the dynamical system (De Martino et al., 2014; Shinar et al., 2009).

A rich plethora of information can be derived from the analysis of the solutions of the dynamical systems associated to the original CR-CRN and to its mutated forms. For example, feedback effects naturally emerge by comparing the time-courses of the individual protein concentrations or by studying the corresponding time-varying reaction fluxes (Sommariva et al., 2021b). More importantly, local and global effects induced on the network by LoF and GoF mutations can be quantified by computing the relative difference between the protein concentrations at the equilibrium point of the original CR-CRN and that at the equilibrium of the network obtained by applying the proper projection operator(s) (Sommariva et al., 2021a). Additionally, analysis of the sensitivity of protein concentrations at equilibrium with respect to the values of the kinetic parameters of the dynamical model may help in identifying the specific reaction or subnetwork mostly affected by each genetic alteration (Biddau et al., 2023).

The proposed model has been used to simulate the functional alterations induced on the CR-CRN by some of the mutations more commonly found in CRC, including the GoF of PI3K, KRAS, and BRAF, and the LoF of PTEN, AKT, and TGF*β*RII (Sommariva et al., 2021a,b; Biddau et al., 2023), and a first attempt in modelling the action of Dabrafenib, a drug targeting BRAF, has been performed by (Sommariva et al., 2021b). The obtained results have been extensively compared with results previously published in literature. However, in all those studies only mutations resulting in a complete LoF or in the highest possible value of GoF of the corresponding protein have been considered. In the present paper we applied the proposed CR-CRN in two novel scenarios: (i) the simulation of global and local effects of different genetic mutations resulting in different levels of alteration of the functional activity of the corresponding proteins; (ii) the in-silico modelling and dosage of a multi-agent targeted therapy. Toward this end, we focused on mutations and drugs involving proteins directly or indirectly related to the mitogen-activated kinase (MAPK), whose overexpression plays an important role in CRC-progression (Fang and Richardson, 2005; Horst et al., 2012; Baharudin et al., 2017; Stefani et al., 2021). More specifically, we quantified the global effects induced on the whole CR-CRN and the local effects induced on the molecules of MAPK by the LoF of PTEN, various levels of GoF of KRAS, and their combination. Finally we investigated the synergies between Dabrafenib and Trametininb, a combination therapy that has demonstrated good results both in terms of progression free survival and response rate (Xie et al., 2020).

## 2 MATERIAL AND METHODS

The equilibrium concentration of the chemical species in both physiological and diseased conditions is interpreted as the asymptotically stable state of the dynamical system

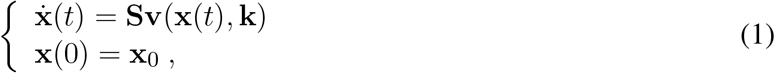

which is obtained by applying the mass action kinetics to the network represented in the molecular interaction map (Kohn, 1999; Pommier et al., 2004; Krogan et al., 2015; Kondratova et al., 2018; Broyde et al., 2021). In equation (1), **x**(*t*) = (*x*_1_(*t*), …, *x*_*n*_(*t*))^*T*^ is the vector whose components are the concentrations of the *n* proteins contained in the network; **k** = (*k*_1_, …, *k*_*r*_)^*T*^ is the vector whose components are the rate constants of the *r* chemical reactions; **S** is the stoichiometric matrix; **v** is the vector of reaction fluxes; and **x**_0_ is the auxiliary initial condition.

The first objective of an analysis of the cancer signalling network performed by means of the dynamical system (1) is the characterization of the equilibrium state of the network as the asymptotic behavior of the solution **x** = **x**(*t*). The main conceptual issue in this respect is that system (1) does not necessarily have a unique equilibrium solution (Feinberg, 1987; Yu and Craciun, 2018; Conradi and Flockerzi, 2012; Conradi and Mincheva, 2014). In order to identify formal assumptions that imply this uniqueness the following process should be considered:

1. Given a solution **x** = **x**(*t*) of (1), a set of *p* semi-positive conservation vectors satisfying *p* conservation laws can be identified, which belongs to the kernel of the transpose of the stoichiometric matrix **S**. The transposed forms of the conservation vectors are used to generate the conservation matrix **N**.
2. The conservation matrix is said weakly elemented if it contains at least one minor equal to the identity matrix of order *p*. If this holds, the solution vector **x** and the conservation matrix **N** can be re-ordered in such a way that the identity matrix acts on the first *p* elements of the re-ordered solution vector.
3. If **N** is weakly elemented, for each initial condition **x**_0_ it is possible to construct *p* hyperplanes in ℝ^*n*^, each one defined by the corresponding conservation law. The stoichiometric compatibility class of **x**_0_ is the intersection of the hyperplanes
4. As said, in general there is no guarantee that the asymptotic solution of (1) is the same for any **x**_0_ ∈ ℝ^*n*^. However, it is possible to formulate a conjecture stating that the asymptotic solution is unique for all initial conditions belonging to the same stoichiometric compatibility class. More precisely, a dynamical system and the corresponding CRN are said to satisfy the global stability condition if such conjecture holds true, i.e. if for every stoichiometric compatibility class there exists a unique globally asymptotically stable equilibrium solution **x**_*e*_.

In order to make a globally stable CRN supportive for the construction of an in-silico model of a colorectal cancer cell, three computational issues should be addressed. First, the dynamical system can be modified in order to implement the presence of single DNA mutations and to compute their impact on the resulting proteomic profile. For example, a LoF mutation, which results in the reduction or even the cancellation of the function of a specific protein, is implemented by projecting the initial concentration values describing the physiological cell onto a new initial state in which the concentrations of the mutated protein and the corresponding compounds are set to zero. On the other hand, a GoF mutation, which enhances the expression of a specific protein, is implemented by setting equal to zero the reaction coefficients corresponding to the equations representing the de-activation reactions for that protein (Sommariva et al., 2021b).

The second issue is concerned with the numerical computation of the (unique) asymptotic solution of (1). Of course, this can be done by applying a numerical method for the solution of the Ordinary Differential Equations (ODEs) contained in the Cauchy’s problem. However, the dynamical computation of this high-dimension system is numerically demanding and can be effectively replaced by numerical optimization. In fact, it can be showed that the stationary solution of the Cauchy’s problem can be determined by an algebraic system, which, in turn, is equivalent to a non-linear root-finding problem. Therefore, in this second approach, iterative schemes can be applied to directly compute the equilibrium state, without the need to approximate the solution of the Cauchy’s problem at each time point (Berra et al., 2022).

Finally, it is important to observe that, on the one hand, system (1) is made of a high number of equations, kinetic parameters, and unknown concentrations; but, on the other hand, that just a smaller subset of such equations is representative of chemical reactions that are actually affected by the cancer somatic mutations. Sensitivity analysis is the technical tool that allows the quantitative assessment of the impact of the kinetic parameters’ uncertainty on the CRN equilibrium state. In its local version, sensitivity analysis is an indicator of the impact that a specific mutation has on the expression of the corresponding protein, and, even more importantly, it is able to identify the sub-networks that are mostly affected by the mutation (Biddau et al., 2023).

## 3 RESULTS

The mathematical model described in (Sommariva et al., 2021a) has been applied by (Sommariva et al., 2021b) to compute modifications in the equilibrium concentrations of the CRC network induced by mutations in a few genes that are rather common in CRC cancerogenesis. In the present paper we focused on a quantitative analysis of the impact of mutations in KRAS, a widely expressed GTP/GDP-binding MAPK protein, whose mutated version is found in more than 30% of CRCs. Specifically, in this section we

- Computed the impact of complete and partial mutations of KRAS on the global proteomic profile of the CRC network.
- Compared the modifications induced by the mutate KRAS on the equilibrium with respect to the ones induced by PTEN, a dual protein/lipid phosphatase that triggers the PI3K/PTEN/AKT signalling pathway.
- Investigated to what extent a mutated KRAS impacts the expression of specific proteins in MAPK.
- Studied both local and global effects of the combination of mutations in KRAS and PTEN.

Then, in the next section, the MAPK pathway will be studied as a molecular target for two inhibitors of BRAF and MEK, respectively.

### 3.1 Global effects induced by mutations in colorectal cancer

Mutations of KRAS are very common in CRC and belong to the pathway of the main sequence k-Ras/Raf/MEK/ERK. In particular, a GoF of KRAS is realized by a modification of the stoichiometric matrix in (1), which, from a chemical viewpoint, corresponds to removing from the CRN all reactions involved in the de-activation of that protein. A study of global effects of the GoF of KRAS on the proteomic profile of a colorectal cancer cell has been studied by (Sommariva et al., 2021b) via the computation of

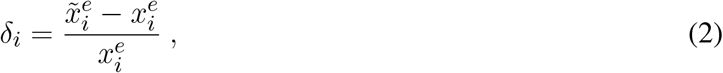

where 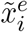 and 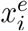 are the mutated and the physiological equilibrium, respectively. Since *δ*_*i*_ and the difference 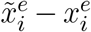 have the same sign, the concentration of the *i*−th protein in the mutated network is either increased if *δ*_*i*_ *>* 0 or reduced if *δ*_*i*_ *<* 0. In particular, a value of *δ*_*i*_ equal to − 1 means that the function of the *I* − protein is completely stopped. In more general terms, the value of *δ*_*i*_ quantifies the relative change of the protein concentration, normalized by its value in the physiological network, and thus enables identifying which proteins are more sensitive to the considered mutation.

Using this same technique, here we studied the consequence of partial GoF mutations of KRAS, i.e. of a common proportional reduction of the values of the rates of those reactions that model the inhibition of the active forms of k-Ras. Precisely, in the top panel of Figure 1 the three profiles correspond to values of the rates set to 0%, 30%, and 60% of the corresponding physiological values, while the second panel from the top shows how the whole network is affected by a complete mutation of KRAS. In the third panel we compared this latter profile with the one associated to another frequent mutation in CRC, i.e. the LoF of PTEN, which belongs to the distinct PI3K/PTEN/AKT pathway. Finally, in the bottom panel we studied the effect of the combination of the two mutations on the CRN global equilibrium.

**Figure 1.**
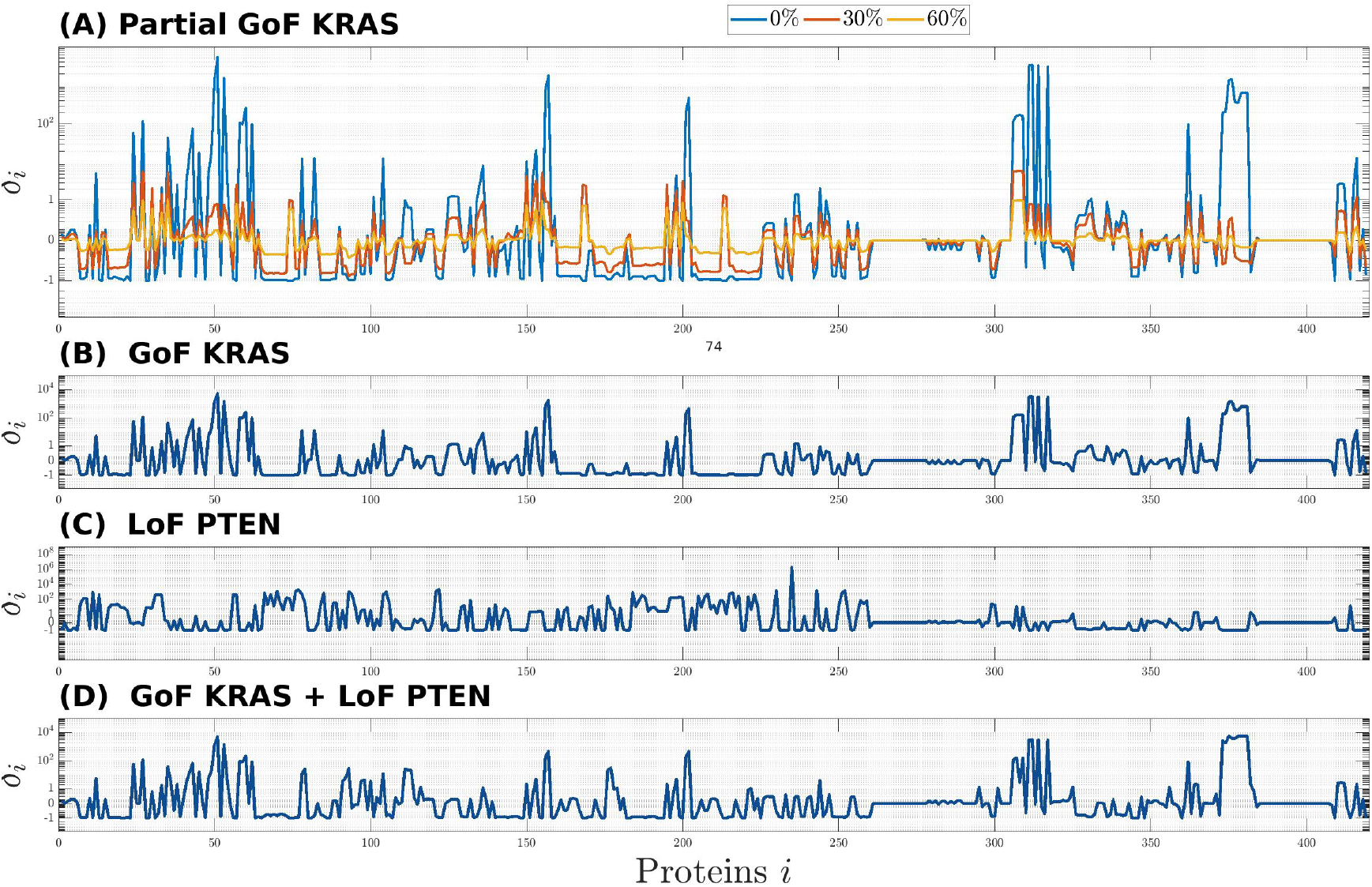
Global effects induced on the CR-CRN by the GoF of KRAS **(A)**-**(B)**, the Lof of PTEN **(C)**, and their combination **(D)**. The effects on each protein, *i* = 1, …, *n*, of the CR-CRN is quantified by the relative difference *δ*_*i*_ between the concentration at equilibrium of the physiological and the mutated networks. Panel **(A)**: results for three mutations corresponding to different levels of GoF of KRAS, obtained by setting the rate constants of the reactions inhibiting the active form of k-Ras to 0%, 30%, and 60% of their physiological values. Panels **(B)** and **(C)**: global effects of the complete GoF of KRAS and LoF of PTEN, respectively. Panel **(D)**: global effects induced by the combination of the two complete mutations.

The results of Figure 1 show that the whole network is affected by the mutations of KRAS, PTEN, and their combination; that, in particular, significant changes in concentration may involve proteins far from the mutated ones in the graph of the CRN; and that the global equilibrium profile changes smoothly with respect to a smooth variation of the reactions’ rate. Further, on the one hand, the higher impact of the GoF of KRAS, which is visible by comparing the second, third, and fourth panels, is possibly related to the fact that this protein is upstream in the global network. On the other hand, the same three panels seem to show that no intuitive superposition principle applies to the profile associated to the combination of mutations. This is probably a consequence of the fact that the effects of the combination follow from the solution of a combined non-linear problem with appropriate initial conditions. Further, the profile of *δ*_*i*_ in panel **(A)** implies a growing difference from the physiological equilibrium, as the values of the mutated coefficients tend to zero, that is, as the mutation is intensified.

With reference to global results concerning the mutation of KRAS, Table 1 shows the list of the 10 molecular species exhibiting the highest variations between physiological and mutated equilibrium values. With the only exception of CDC25C, all of them belong to the mitogen-activated kinase (MAPK) signalling pathway, consisting of the main sequence k-Ras/Raf/MEK/ERK, while Pase3 is a phosphatase acting on phosphorylated ERK proteins. This result, ultimately obtained from the simulation of the mutation of KRAS, agrees with well known aspects of the physiology of the MAPK signaling pathway. Essentially, the activated form of k-Ras is responsible for the transduction of signals, received at the cell surface, to the inside of the cell; this operation is crucial for cell proliferation, growth and differentiation (Guo et al., 2020; Lavoie et al., 2020; Morkel et al., 2015; Pappalardo et al., 2016; Porru et al., 2018). Under the GoF mutation the molecules of k-Ras persist in their active form, which implies an aberrant activation of downstream effectors as Raf, MEK, ERK, and leads to a malignant behavior of the cell. Thus the whole signaling transduction pathway MAPK is uncontrollably triggered by the alteration of KRAS and may lead to out of control cell proliferation. The protein ERK exhibits the maximum difference between mutated and physiological equilibrium values; further, its active form p-p-ERK, which occupies the third place in the list of Table 1, is a well known “master regulator of cell behavior, life and fate” (Lavoie et al., 2020), being deeply involved in cellular responses as cell proliferation, survival, growth, metabolism, migration and differentiation (Guo et al., 2020; Lavoie et al., 2020; Sugiura et al., 2021). This makes the MAPK path a very natural target of drugs which contrast the negative effects of the mutation of KRAS. In particular, the response of active ERK to the delivery of drugs has received special attention (Hamis et al., 2021; Lavoie et al., 2020; Pappalardo et al., 2016; Santini et al., 2019). Finally, we observe that the high value of the relative difference *δ*_*i*_ of the protein p-p-ERK is due to the smallness of the related physiological equilibrium value.

**Table 1.** Proteins showing the most significant variation in concentration (reported in decreasing order) when the network is affected by the complete GoF of KRAS. Such change is quantified by the difference between the concentration values at equilibrium in the physiological status and after the mutation (second column), and by the value of the corresponding *δ*_*i*_ (third column).

| i-th protein | $\vert \tilde{x}_i^e - x_i^e\vert $ | $\delta_i$ |
| --- | --- | --- |
| ERK | 179.31 | -0.93 |
| MEK | 128.35 | -0.77 |
| p-p-ERK | 91.95 | 5365.73 |
| k-Ras_GDP | 82.39 | -0.98 |
| k-Ras_GTP | 69.69 | 118.38 |
| Pase3 | 67.31 | -0.71 |
| p-p-MEK | 62.84 | 76.78 |
| p-p-ERK_Pase3 | 41.94 | 1550.88 |
| p-MEK | 37.47 | 3.21 |
| CDC25C | 36.96 | -1.00 |

As a final comment concerning Figure 1 and Table 1, we remark that all these global results follow from the simulated behavior of the equilibrium solutions of a system of ODEs. Relevant knowledge from cell biology and biochemistry has been applied in the formulation of the basic physiological model of the network and in the description of mutations (Tortolina et al., 2015). Explicit information on possible biological consequences can be recovered when looking at the local scale.

### 3.2 Local effects induced by mutations in colorectal cancer

An analysis of local effects of mutations may lead to a more direct connection with the physiology of the signaling network. A few items have already been examined in (Sommariva et al., 2021b), mainly focused on the study of the effects of various single-gene mutations on the concentration values of p53. Here we present a further set of complementary results on the MAPK pathway.

Figure 2 is devoted to a quantitative analysis of the changes of the MAPK cascade induced by mutations. We have dedicated a panel to each of the molecular species k-Ras, Raf, MEK, ERK, and to the corresponding active forms k-Ras_−_GTP, p-Raf, p-p-MEK, p-p-ERK. The histograms inside each panel provide the concentration values at equilibrium in the following conditions: physiological (purple), mutated by GoF of KRAS (blue), mutated by LoF of PTEN (green), combination of the last two (yellow). Note that the range of the *y*-axis depends on the panel. The consequences of the mutations outlined by the histograms agree with a number of remarks on the physiology of the MAPK cascade that are scattered over the literature (Morkel et al., 2015; Santini et al., 2019; Guo et al., 2020; Lavoie et al., 2020)

**Figure 2.**
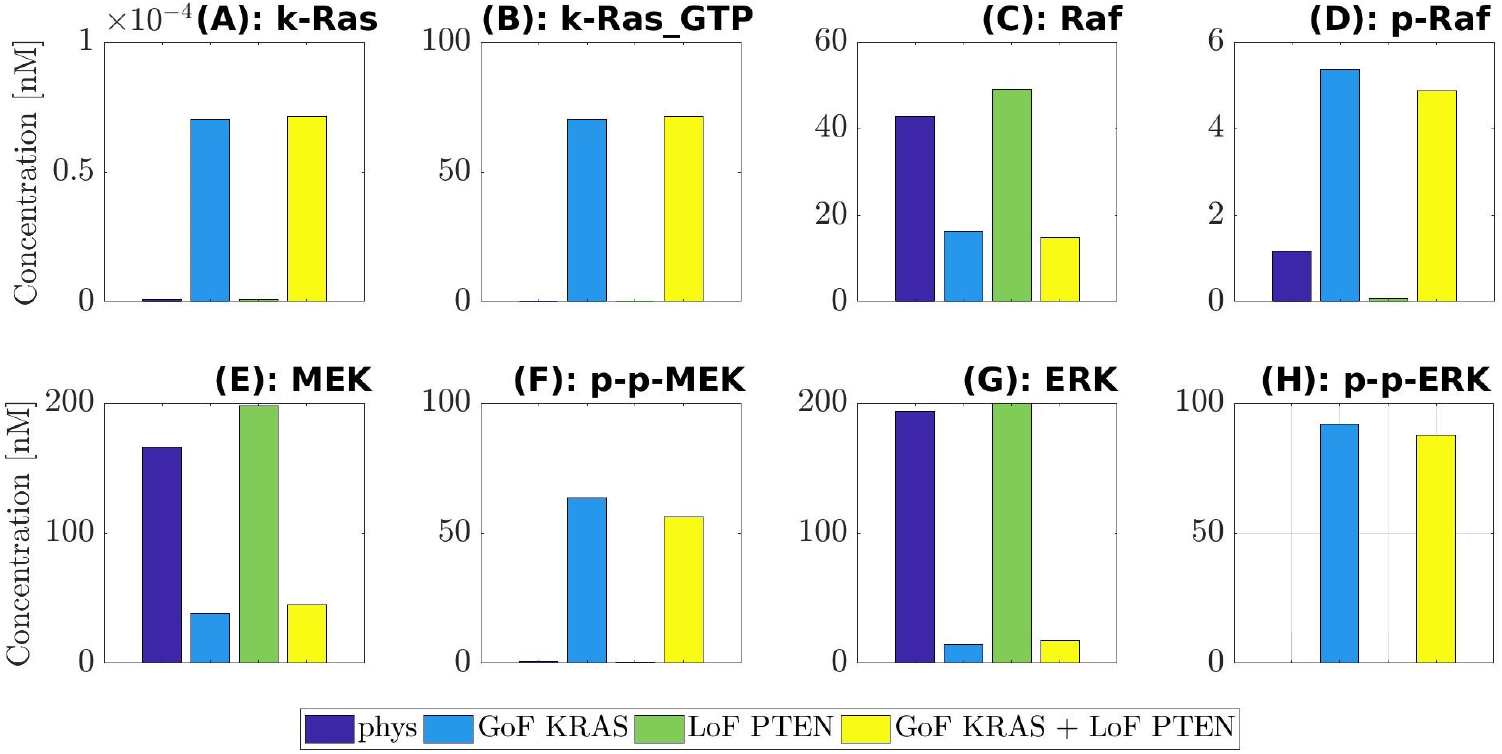
Values of the concentration at equilibrium of the four species belonging to MAPK pathway (k-Ras, Raf, MEK, ERK) and of their active forms (Ras GTP, p-Raf, p-p-MEK, p-p-ERK). The computed equilibria are related to the physiological network (purple) and to the same one but affected by three different mutations: GoF of KRAS (blue), LoF of PTEN (green) and the concurrence of the two (yellow). For ease of visualization a different scale is used on the y-axis of each label.

For example, the first two columns of each panel compare physiological equilibria (purple) with those mutated by the GoF of KRAS (blue). We observe that the mutated value of k-Ras remains rather small, although increased as a consequence of the mutation. As an expected result of the GoF, the value of the active form k-Ras_−_GTP is raised to about 70 nM. Also, the physiological concentration of Raf is almost halved, while the mutated value of the active form is about five times the physiological one. Similarly, the inactive form of MEK is heavily decreased by the mutation, while the active form p-p-MEK is highly augmented; the same remark holds for ERK and p-p-ERK. To summarize, the concentrations of the inactive forms of the key elements of the MAPK pathway are reduced by the mutation of KRAS (with the only exception of k-Ras), while the concentrations of the active forms are heavily enhanced. Consequently, the whole path is abnormally active.

Consideration on the first (purple) and the third (green) columns provides the impact of the LoF of PTEN on the MAPK pathway. It is found that the equilibrium changes of k-Ras, k-Ras_−_GTP, p-p-MEK, ERK, p-p-ERK can be overlooked, while small changes can be seen in Raf, p-Raf, MEK. Since the mutated active forms of the basic elements of MAPK are almost vanishing, we may conclude that there are no sensible effects of the LoF of PTEN on MAPK. An inspection of the fourth columns (yellow) in comparison with the second ones (blue) shows that the combination GoF of KRAS + LoF of PTEN produces essentially the same effects as the mutation of KRAS alone. This is a further indication that the MAPK path is not sensitive to LoF mutations of PTEN.

The marked increase of the concentration of the active form of ERK under the GoF of KRAS, with the consistent reduction of the inactive ERK, leads in particular to abnormal cell proliferation (Hamis et al., 2021; Lavoie et al., 2020), as already observed while commenting Table 1. Further, it is known (Sugiura et al., 2021) that ERK promotes the apoptosis of the cell, and that this property is inhibited by the activation of the species itself. Looking at the histograms, we can observe that the quantity at equilibrium of p-p-ERK grows steeply under the mutation of KRAS, while the concentration level of ERK decreases significantly. It follows that ERK’s anti-proliferative effects are missing and tumor cell death is not induced anymore, confirming the danger of such mutation.

### 3.3 Drugs and drug combination targeting the MAPK pathway

The main objective of this section is to simulate the impact of drugs having the MAPK pathway as their target. Our investigation follows the previous analysis of mutations, by considering first the global effects on the whole network, and next the local effects of the MAPK pathway. As to the global effects, we examined the equilibrium states; as to the local effects, we focused on the analysis of the time course of the activated protein p-p-ERK, which is the key factor to assess the main effects arising from the MAPK pathway. In the present approach, the optimal concentration of a drug is determined by the degree of compliance between the equilibrium features of the model modified by the drug, and the corresponding features of the original physiologic model.

To describe the action of a given drug, we enlarge the set of the chemical reactions of the CRC CRN, and the corresponding dynamical system, in order to account for the reactions between the drug and the target molecules. As an immediate consequence, the drug and the associated composites are regarded as additional unknowns. Borrowing from a rather well established literature, we have simulated the action of two drugs: Dabrafenib (DBF) and Trametininb (TMT), with target MEK. As far as reactions are concerned, DBF is modeled as a competitive inhibitor of Raf, while TMT is an allosteric inhibitor of MEK (Hamis et al., 2021; Morkel et al., 2015; Puszkiel et al., 2019; Sommariva et al., 2021b) (more details are given in the Appendix). The key parameters of the enlarged system are represented by the initial values of the drug concentrations, say *c*_*D*_ for DBF, and *c*_*T*_ for TMT, although also the additional rate constants have to be fixed; the initial values *c*_*D*_ and *c*_*T*_ denote also the total amounts of drug available for the network.

For each inhibitor we examined global and local effects; also, effects of combinations of DBF and TMT are illustrated. In the present approach we considered the mutated equilibrium state as the initial state of the dynamical system modified by the addition of a drug, and we determined the resulting new equilibrium state 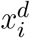. In particular, we find the most appropriate concentrations, *c*_*D*_ and/or *c*_*T*_, to obtain an equilibrium 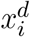 closest to the physiologic equilibrium 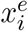. Figure 3 provides a synthetic description of the changes induced by the action of DBF on the network subject to a GoF mutation of KRAS. This figure has been obtained through the following steps.

**Figure 3.**
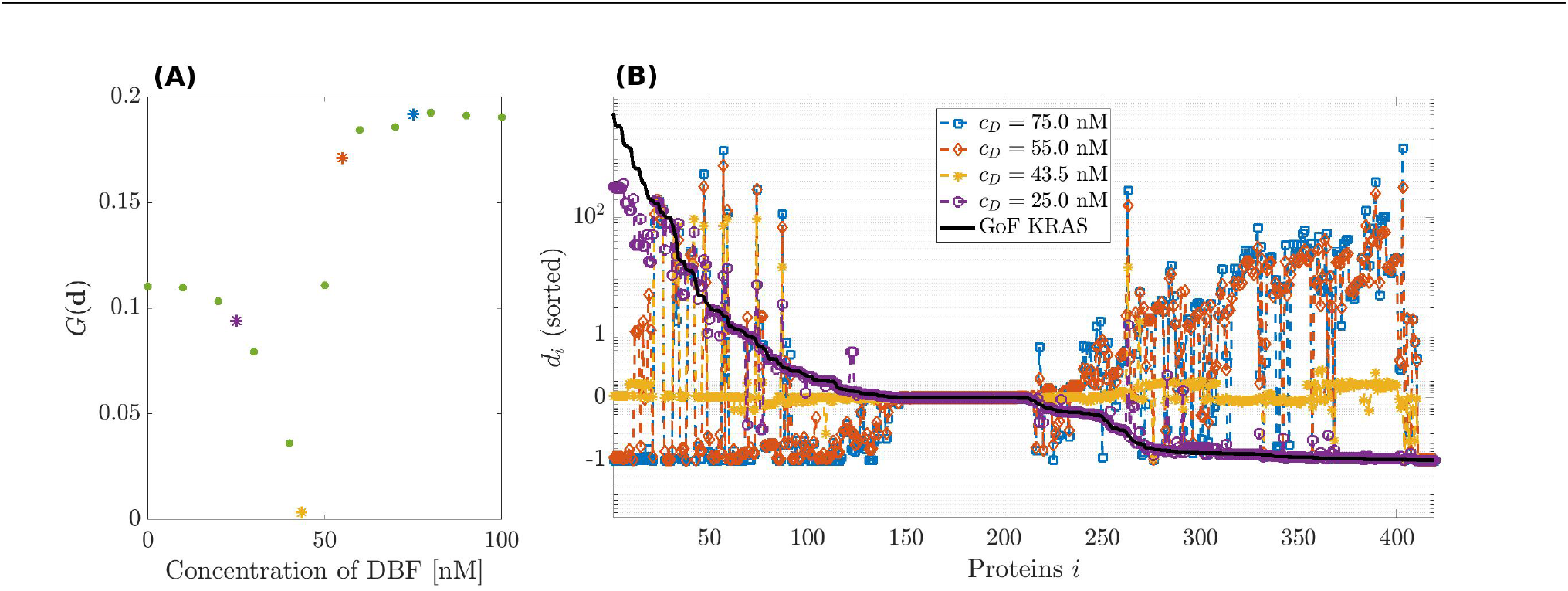
Dosage of Dabrafenib (DFB). Panel **(A)**: modified geometric mean *G* computed on vector (**d**) = (*d*_*i*_)_*i*_ of the relative differences between the proteins’ concentration at the equilibrium of the physiological network and in the one obtained by simulating the action of DBF against the GoF mutation of KRAS. Several values for the initial concentration *c*_*D*_ of DBF have been tested and the colored marks corresponds to those further explored in the right panel. Panel **(B)**: Relative difference *d*_*i*_ as function of the proteins quantifying the effect of the GoF mutation of KRAS (black line) and the effect of four initial values of DBF concentration (colored lines).

- Consider the augmented system formed by the mutated dynamical system, enlarged by addition of the reactions expressing the action of DBF.
- Consider the mutated equilibrium state 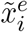 and the initial value for the drug concentration *c*_*D*_ as the initial values for the dynamics of the augmented system.
- Determine the equilibrium state 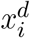 of the augmented system.
- Compare 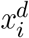 with the physiological equilibrium 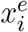 by considering the corresponding relative difference of the concentrations, which here is denoted as

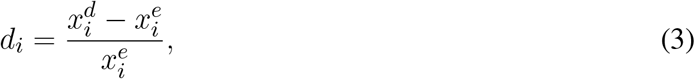

for convenience.
- Investigate the dependence of *d*_*i*_ on *c*_*D*_, and plot the results.

Clearly, the procedure also applies to the description of changes induced by TMT, with initial value *c*_*T*_, and to the combination of the two drugs. The underlying idea is that the drug works well at the concentration considered if the quantities *d*_*i*_ are small, that is, if the equilibrium state reached under the action of the drug is close to the physiological (healthy) state.

In order to determine which is the best quantity of drug to be administered the information in vector **d** = (*d*)_*i*_ is summarized in a unique numeric value through the indicator

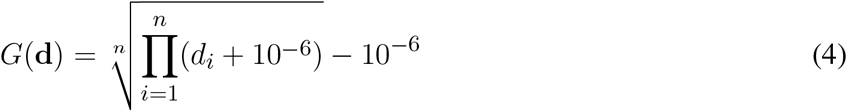

that consists in a modified geometric mean allowing the vector **d** to contain elements equal to zero (De La Cruz and Kreft, 2018). Such index is exploited in Figures 3, 4, 5 for showing the performance of single drugs DBF and TMT and their combination in dependence of their initial concentration values inside the network. The lower the value of *G*(**d**), the closer the equilibrium of the network including the drug(s) is to the psychological equilibrium. Therefore, the optimal initial concentration for each therapy could be determined by minimizing *G*(**d**).

**Figure 4.**
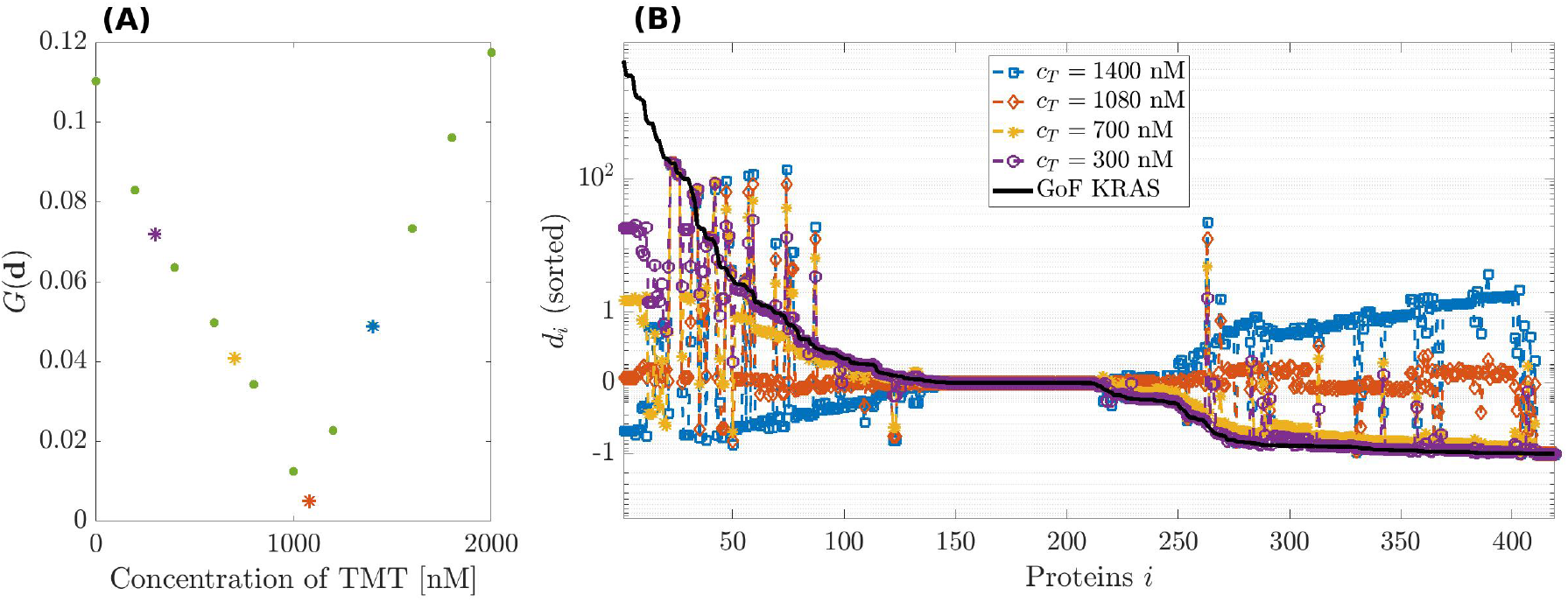
Dosage of Trametinib (TMT). Panels **(A)** and **(B)** have been designed as in Figure 3 but considering TMT instead of DBF.

**Figure 5.**
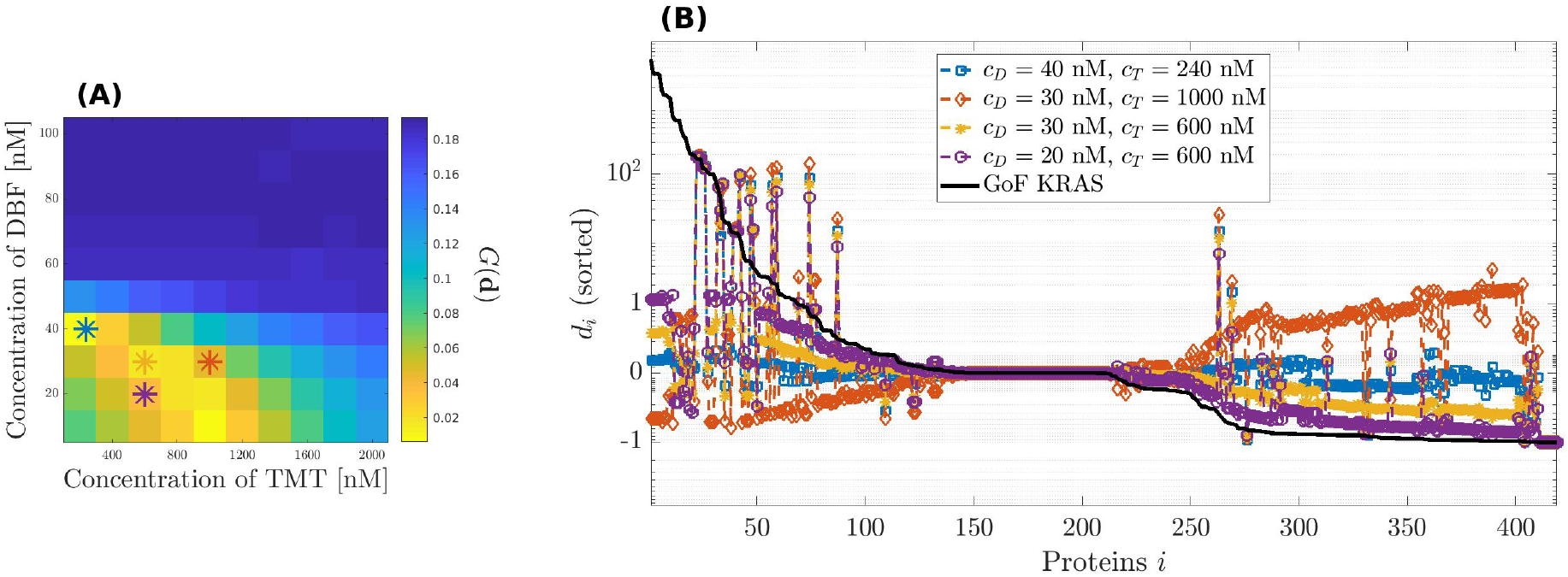
Dosage of the combination therapy including Dabrafenib (DBF) and Trametinib (TMT). Panel **(A)**: 2-D images representing the modified geometric mean *G* of the vector ***d*** = (*d*_*i*_)_*i*_ of the relative differences between the proteins’ concentration at the equilibrium of the physiological network and in the network obtained by simulating the concurrent action of DBF and TMT against the GoF mutation of KRAS. Several pairs of values (*c*_*D*_, *c*_*T*_) for the initial concentrations of the two drugs have been tested and the colored marks correspond to those further explored in the right panel. Panel **(B)**: Relative difference *δ*_*i*_ as function of the proteins quantifying the effect of the GoF mutation of KRAS (black line) and the effect of four pairs of initial concentration values of DBF, *c*_*D*_, and TMT, *c*_*T*_ (colored lines).

Figure 3, panel **(A)** shows the plot of the modified geometric mean *G*(***d***) as a function of the initial concentration of DBF *c*_*D*_, which varies in the interval [0, 100] nM. Therefore, the plot provides an estimate of the drug potential of restoring the healthy state of a cell. The minimum of *G* corresponds to the initial drug concentration that assures the best healing effect on the mutated network. Figure 3 panel **(B)** shows a few complete profiles of the relative differences *d*_*i*_. The black line, which has been inserted for ease of comparison, represents the relative difference between the concentrations at equilibrium under a GoF of KRAS and the physiological equilibrium; notice that the values of *δ*_*i*_ are sorted in decreasing order, to give evidence to the changes induced by the drug. The different colors of the other lines correspond to the profiles of *d*_*i*_ determined by different values of *c*_*D*_.

The best choice of *c*_*D*_ corresponds to the line closest to the horizontal axis, i.e., *c*_*D*_ = 43.5 nM. This is in essential agreement with the result of panel **(A)**. Notice that the initial values of the concentration of DBF and the values of the rate constants of the related reaction have been changed with respect to those used in (Sommariva et al., 2021b), as observed in the Appendix; this explains the slight difference with the optimal value of the initial concentration found in that paper.

In correspondence with the value *c*_*D*_ = 43.5 nM, the concentrations of most proteins involved in network are very close to the values at the physiological equilibrium. We point out a few exception: Raf, whose concentration is reduced in that its function has been inhibited by the drug; a group of complexes that involve the activated form of k-Ras, that is still overexpressed; the complexes that are products of the reactions removed to simulate the GoF of KRAS, whose function is thus stopped.

Figure 4 describes the same analysis, this time performed in the case of the effects of TMT on the GoF of KRAS. In this case, the initial concentration values are allowed to vary over a rather large interval, from 200 to 2000 nM, the system being poorly sensitive to variations of *c*_*T*_. A minimum of *G*(**d**) is found at *c*_*T*_ = 1080 nM. Figure 4, panel **(B)**, agrees with the previous result.

Finally, Figure 5 analyzes the effects on the relative error *d*_*i*_ of various combination therapies involving DBF and TMT. The heatmap of panel **(A)** shows the response to a set of combination therapies. On the horizontal axis, the concentration of TMT varies from 0 to 2000 nM; on the vertical axis, the concentration of DBF describes the interval from 0 to 100 nM. The value of *G*(**d**) in each square is inferred by comparison with the colors of the vertical bar. The best result is obtained for *c*_*D*_ ≈ 40 nM and *c*_*T*_ ≈ 200 nM. Thus, a combination of smaller doses of both drugs produces almost the same effects of either a single infusion of DBF, or TMT, delivered at much higher dose.

Panel **(B)** shows four examples of *d*_*i*_ distribution, corresponding to different choices of the initial conditions *c*_*D*_ and *c*_*T*_ for DBF and TMT. As in the previous analogous representations of the distributions associated with drugs, we have reported by a black line the relative difference *δ*_*i*_ between equilibrium values of the network subject to the GoF of KRAS and the physiological values. Comparison of the results shows that there is a significant convergence between the conclusions drawn from panels **(A)** and **(B)** as to the most convenient combination.

The last part of this section is devoted to local considerations on the time course of p-p-ERK. For the ease of reference, we recall that ERK is an elemental conserved variable, according to (Sommariva et al., 2021a). Thus, we denote by ERK_tot_ the corresponding conserved value, and we call activated fraction of ERK the ratio p-p-ERK(*t*) / ERK_tot_. Fig (6) shows the time course of the activated fraction of ERK under the action of DBF in panel **(A)**, TMT in panel **(B)**, and the combination DBF + TMT in panel **(C)**. The initial conditions for the drugs are chosen as *c*_*D*_ ∈ {12.5, 25, 37.5, 50} (nM), *c*_*T*_ ∈ {50, 100, 150, 200} (nM), and the same values are considered for the drug combinations.

**Figure 6.**
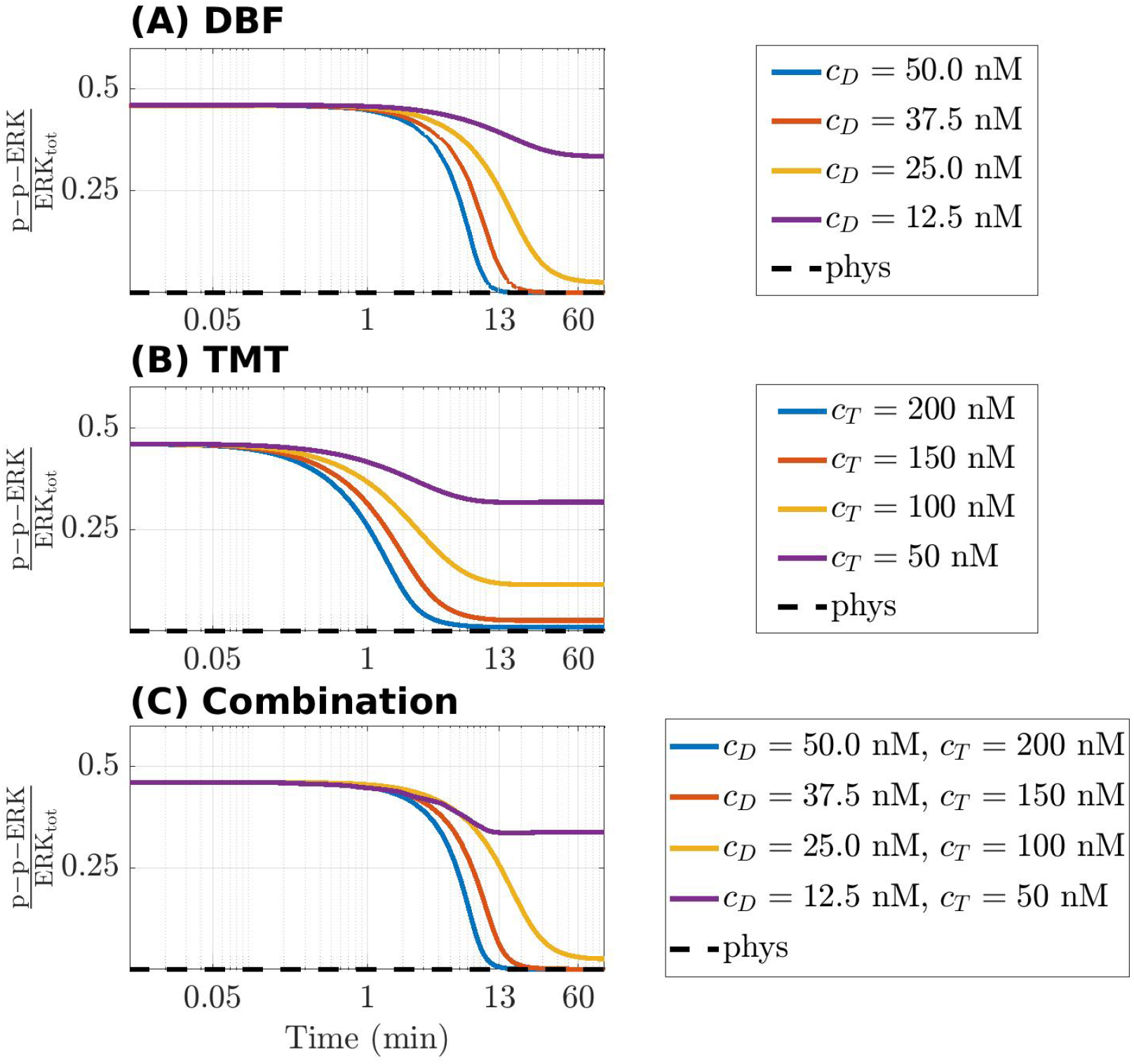
Time-varying behaviour of the concentration of p-p-ERK, active form of ERK, in the CR-CRN affected by GoF of KRAS under the action of DBF **(A)**, TMT **(B)** and their combination **(C)**. In all panels, the activated fraction of ERK is represented. The black dashed line indicates the value of active ERK at equilibrium in physiological conditions, while colored lines refer to four different therapeutic dosage as specified in the legend of each panel where *c*_*D*_ and *c*_*T*_ indicate the initial concentration of DBF and TMT, respectively. A logarithmic scale is used on the time axis.

The graphs of the three panels have a rather similar behavior. A common feature is that the huge amount of the activated fraction of ERK decreases with time, because of the indirect action of the drugs that target Raf or MEK or both. In the case that only small quantities of delivered drug are available, each panel contains a curve showing an initial slight decrease of the ratio, until it tends to a non vanishing constant value for growing *t*. On the contrary, the curves associated with higher amounts of drug show an almost horizontal behavior for a brief initial time interval, after which they decrease very steeply until a rather small value of the activated fraction of ERK is found. Actually, the almost stationary low value is reached in about 13 minutes or more for DBF and TMT+DBF, and in about 6 minutes under the action of TMT alone, perhaps because the target MEK of TMT is closer to ERK in the network topology.

## 4 DISCUSSION

A mathematical model has been recently introduced (Tortolina et al., 2015; Sommariva et al., 2021a) that simulates the behaviour of a CRN describing the information flow inside a CRC cell at the G1/S transition point; furthermore, general procedures to change the model in order to account for GoF and LoF mutations have been proposed and investigated (Sommariva et al., 2021b). In other words, the transition from a healthy to a mutated, cancerous, signaling network has received an appropriate mathematical formulation.

In the present study the mathematical model has been applied in order to analyze the reaction of the network to a GoF mutation of KRAS, a LoF of PTEN, the combination of the two mutations, and a partial GoF mutation of KRAS. The interest towards mutations of KRAS and PTEN comes from the observation that they have been found in approximately 40% and 34 % of all CRC cases, respectively (Zhu et al., 2021; Salvatore et al., 2019). A global analysis of pertinent equilibrium states has pointed out the MAPK pathway, with its k-Ras/Raf/MEK/ERK cascade, as the target of the main changes of the network, induced by the mutation of KRAS. A corresponding local analysis has provided a suggestive graphical representation of quantitative aspects of the products of activation/inactivation reactions. In particular, the mathematical analysis identifies the activated form of ERK as the focus of changes, in parallel with well known results coming from the biologic side (Guo et al., 2020; Lavoie et al., 2020; Sugiura et al., 2021).

Thus, the species of the MAPK pathway represent the natural target of drugs that tend to contrast the negative consequences of k-Raf mutations. Here we have examined the response of the mutated network to administration of DBF and TMT, with respective targets Raf and MEK; more precisely, DBF is modeled as a competitive inhibitor of Raf, and TMT as an allosteric inhibitor of MEK. As a further related development, the effects of combination of the two drugs at variable doses have also been simulated. In fact, it is well known that drug combinations may counterbalance, e.g., the onset of drug-resistant tumour subclones (Hamis et al., 2021; Morkel et al., 2015; Santini et al., 2019).

The main novelty of the mathematical scheme for the simulation of drug effects is given by application of three different models, namely, the model for the healthy, the mutated, and the drug loaded network. Here, the mutated network has played a fundamental role. In our approach, we have considered the equilibrium states pertaining to each model. The mutated equilibrium has provided the initial value of the drug loaded model, leading to the associated equilibrium. This last, in turn, has been compared with the healthy equilibrium. A drug, or a drug combination, has been regarded as effective if its drug loaded equilibrium is close to healthy equilibrium, which means that the relative difference *d*_*i*_ is globally small.

Next, a global index has been introduced, dependent on *d*_*i*_, which selects the amount of drug generating the closest model to physiologic equilibrium. The procedure has been applied to assess the optimal concentrations of DBF and TMT. The results have been confirmed by graphical representations of the distribution of the relative differences between drug loaded and physiological equilibrium. Similarly, an optimal drug combination of DBF and TMT has been assessed on the basis of the values of the global index that have been reported on a heatmap. Again, the optimal choice has been validated by the representation of the distributions of relative differences.

At the local level of analysis, we have examined the time course of the ratio p-p-ERK(*t*)/ERK_tot_, representing a fractional measure of the activated ERK. Unlike recent computational approaches as (Hamis et al., 2021; Pappalardo et al., 2016), which, however, consider mutations of Ras, we have obtained a realistic behavior of the activated fraction of ERK, whereby the ratio decreases with time, because of drug action. In our opinion this result follows ultimately from the choice of the non-vanishing initial value of p-p-ERK, coincident with that of the mutated equilibrium, and the global effects of the network, incorporating also, e.g., feedback effects.

To further ascertain the realistic behavior of active fraction of ERK that follows from our approach, we have investigated the changes induced under the assumption that DBF is subject to degradation, while TMT is absent. Precisely, we have considered for DBF a degradation rate equal to 5.79·10^−6^ s^−1^ (Anderson et al., 2019), while all other conditions of the model have been left unaltered. Fig 7 shows the results: the active local fraction of ERK shows an initial value of about 0.5, which is reduced to a very small value in the first hour; the latter is maintained for a rather long time interval, until it reaches again the value 0.5, following the decrease of the drug concentration caused by degradation.

**Figure 7.**
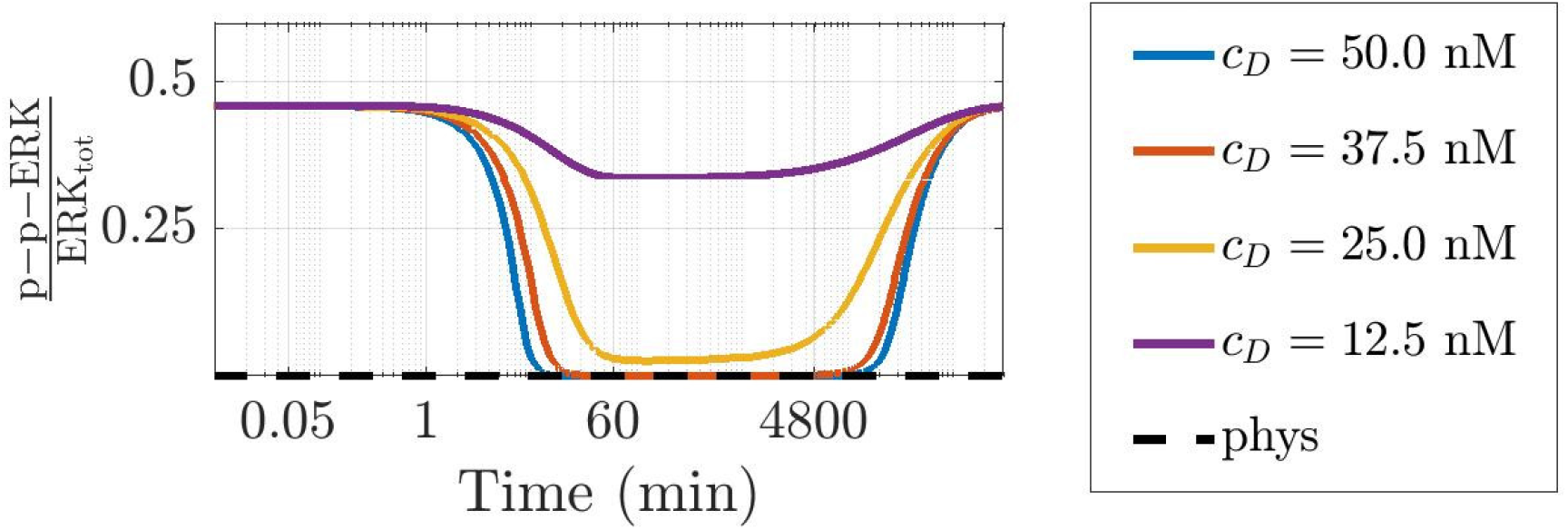
Time-varying behaviour of the concentration of p-p-ERK, active form of ERK, in the CR-CRN affected by GoF of KRAS under the action of DBF when also a reaction modeling drug degradation is included in the model. The figure has been designed as panel **(A)** of Figure 6. A logarithmic scale is used on the time axis; *c*_*D*_ indicated DBF initial concentration.

## 5 CONCLUSIONS

In this work we have first reviewed the most fundamental aspects of a recently proposed CRN which simulates the behavior of the signaling network inside a colorectal mutated cell (Sommariva et al., 2021a,b). We have described the general context where the CRN is applied, the basic principles underlying the development of a system of ODEs representing the chemical reactions of the network, and we have reviewed the most fundamental properties of the ODEs applied in the simulations.

Next, we have developed new applications concerning comparison between a physiological (healthy) network and a few similar networks resulting from GoF mutations of KRAS, and LoF mutations of PTEN, frequently observed in CRC. Also, drug loaded networks associated with either single or combined targeted drugs have been investigated and compared. The results obtained have been validated using literature data.

The basic novelty of our approach is given by the interaction between global and local aspects in the treatment of mutations and the action of drugs. The most interesting example is concerned with the abnormal value of activated ERK (p-p-ERK) resulting from a GoF of KRAS; unlike other approaches (Hamis et al., 2021; Pappalardo et al., 2016) it is found from the simulations that activated ERK is considerably reduced by administration of DBF, a Raf targeting drug. The local time course of the complex p-p-ERK, which is of fundamental interest because of its biologic consequences (Hamis et al., 2021; Guo et al., 2020; Lavoie et al., 2020; Sugiura et al., 2021), is obtained through the use of three different, global, and interconnected networks (physiological, mutated, drug loaded) to assess the general framework which provides both the system of ODEs to be solved, and the required initial conditions.

The methods that have been proposed in this paper may be applied to the prediction of quantitative effects of targeted drugs, and to the optimization of combination therapies for the mutated cell of other cancer types, under rather general conditions.

In our analysis we have assumed that the parameters of the model are fixed and given, and that a unique equilibrium state exists for every stoichiometric surface. These points need for further investigation. For example, a sensitivity analysis is required to first identify those parameters that are most influent on the equilibrium values, and then to design proper biological experiment to refine their values.

Also, the basic model may require adjustments in order to account for specific effects as the natural process of degradation of drugs and other proteins, or changes of the interactions between proteins, possibly occurring as an answer of the network to modifications induced by drugs.

Finally, we should model glucose metabolism and cell apoptosis. In a sense, our approach to drug action is almost opposite to that of inducing cell apoptosis, in that our main aim has been to restore a (nearly) healthy state, instead of triggering cell death.

## 6 APPENDIX

DBF has been modeled as an inhibitor of Raf activation (Hamis et al., 2021; Morkel et al., 2015; Sommariva et al., 2021b). Thus, we have considered the CR-CRN that takes into account the GoF mutation of KRAS, modified by addition of the reversible reaction (Sommariva et al., 2021b)

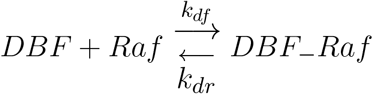

where DBF_−_Raf is the inactive drug target complex. Here the competition is between the inhibitor DBF and Ras GTP, both binding the same site of Raf. The values of the rates are taken from (Hamis et al., 2021) as *k*_*df*_ = 0.106·10^−3^ nM^−1^s^−1^, *k*_*dr*_ = 0.593·10^−4^s^−1^. Notice that they are different from those considered in (Sommariva et al., 2021b). In the search for the steady state under the action of the drug, the initial values of the protein concentrations have been set equal to the equilibrium values of the network subject to the GoF mutation of KRAS. Taking inspiration both from (Hamis et al., 2021) and from (Sommariva et al., 2021b), the initial values of the drug concentration have been extracted from the interval [0, 100] nM.

TMT has been modelled as an allosteric inhibitor of p-p-MEK (Hamis et al., 2021; Morkel et al., 2015), which in turn acts as an enzyme for the double phosphorylation of the substrate ERK. Indeed, ERK is activated by the reaction with p-p-MEK which binds at a certain site, while TMT binds at a different site of the p-p-MEK molecule. Thus the complex TMT_−_p-p-MEK may bind ERK to form the inert complex TMT_−_p-p-MEK_−_ERK, which cannot produce a phosphorylated ERK. The action of TMT has been described by adding to the mutated network the reactions

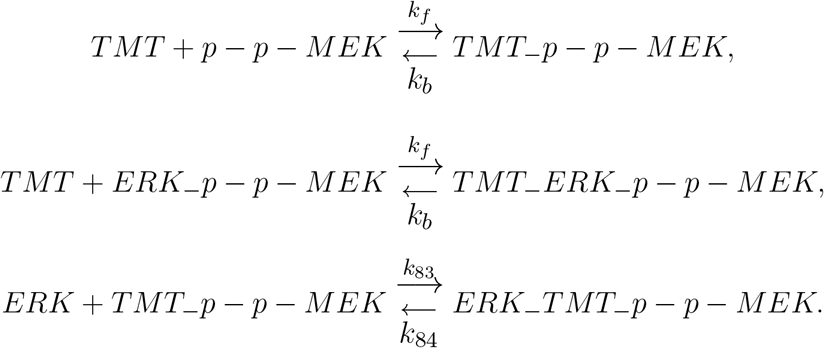

The values of the rate constants *k*_*f*_, *k*_*b*_ have been taken from (Hamis et al., 2021) as *k*_*f*_ = 0.106·10^−3^ nM^−1^s^−1^, *k*_*b*_ = 0.12296·10^−2^ s^−1^, whereas *k*_83_ and *k*_84_ have been taken from (Sommariva et al., 2021a) as *k*_83_ = 0.1·10^−1^ n M^−1^s^−1^ and *k*_84_ = 0.33·10^−2^s^−1^. Following (Hamis et al., 2021), the values of the initial concentration of TMT have been chosen in the interval [0, 2000] nM.

## AUTHOR CONTRIBUTIONS

SS, GC, and MP conceived the paper and designed the computational tests. All authors contributed in optimizing the dynamical model and the algorithms for computing its equilibrium states. SB realized the simulations. SS, MP, GC, and SB wrote the text of the manuscript. All authors validated the results and revised the final text of the manuscript.

## ACKNOWLEDGMENTS

Our colleague Gabriele Zoppoli is kindly acknowledged for discussion on the results of the paper.

## DATA AVAILABILITY STATEMENT

Publicly available datasets were analyzed in this study. This data can be found here: https://github.com/theMIDAgroup/CRC_CRN.

